# Analyzing effect of altitudinal variation in Enzymatic antioxidants of *Coleus forskohlii* from Uttarakhand, India

**DOI:** 10.1101/662528

**Authors:** Pawan Singh Rana, Pooja Saklani

## Abstract

Enzymatic antioxidant activity of the five populations of the medicinal plant *Coleus forskohlii* from five locations of varied altitudes was assayed to analyze the effect of altitude on the enzymatic antioxidant potential. The various enzymes assayed were SOD, CAT, POD, PPO, APX and GR. Highest activity for all the enzymes was observed at higher altitudes. Strong positive correlation was observed between the protein content, enzyme activities and altitude. CAT, POD and GR activity increases significantly with the altitude while SOD was least affected. APX and PPO shows positive correlation. High activity of all these enzymes seems to be to combat the high oxidative stress at higher altitudes. Results of the present study suggest that *Coleus forskohlii* population growing at a higher altitude has higher antioxidant potential than those at lower altitude. Thus, the population of *Coleus forskohlii* from a higher altitude can be used as a source of antioxidants and for commercial propagation.

## Introduction

Environmental factors like temperature, atmospheric pressure and light radiation changes readily with the elevation. At higher altitudes low temperature, enhanced UV radiation and low atmospheric pressure pose serious environmental stresses to the plant growing in the environment (**Barry, 1992**). High light intensities at higher altitudes leads to accumulation of reactive oxygen species (ROS) that can induce protein damage, lipid peroxidation, enzyme inactivation and cell death (**Moller et al., 2007; Latemona et al., 2013;**) To survive such stresses plants growing under these conditions develops some changes in their physiology that includes osmoregulation, hormone level, cell membrane saturation and antioxidant defense (**Huang et al., 2014**). Adaptability is a long term evolutionary process and the plants growing in such environment supposed to have differences from their counterparts growing at a lower altitude. Enzymes forms main component of antioxidant defense system, these include Superoxide dismutase (SOD), Peroxidase (POD), Catalase (CAT), Ascorbate peroxidase (APX), Polyphenol oxidase (PPO) and Glutathione reductase (GR) (**Noctor et al., 1998**). All these enzymes help in maintaining the redox state of plant cells by decomposing H_2_O_2_, superoxide anions and other free radicals (**Cho et al., 2005**). Under stress conditions of higher altitudes, an increase in activity of these enzymes is possible in order to combat the reactive oxygen species produced. Enhanced activity of antioxidant defense in plant under various environmental stresses have been studied by many workers (**Zaefyzadeh et al., 2009; Chen et al., 2010**).

*Coleus forskohlii* is an important medicinal plant grows in wide range of altitudes from 600-2300m (**Chandel and Sharma, 1997**). The plant is well described in Ayurveda for its medicinal properties and is endemic to India (**Valdes et al., 1987; Patil et al., 2001**). It grows wild in the Himalayan region and can be seen easily on the dry barren hills, wasteland and agricultural fields (**Kurian and Sankar, 2007**). *Coleus forskohlii* is widely used in the Indian subcontinent and many countries against various ailments. It is used as emmenagouge, cough expectorant and diuretic in Africa, in treating intestinal and stomach disorders in Brazil and in treatment of cardiac disorders, insomnia, convulsions and respiratory disorders in India (**Valdes et al., 1987; Ammon and Muller, 1985**). *Coleus forskohlii* also claimed to possess antioxidant activities as well (**Khatun et al., 2011**) Thus, considering the medicinal value and wide growth range, the present study was undertaken to study the enzymatic potential of five coleus populations collected from different altitudes.

## Material and Methods

### Plant material

The fresh leaf samples of *Coleus forskohlii* were collected from five selected locations of varying altitudes from Garhwal region of Uttarakhand, India. The leaves were washed with tap water and kept in thermal proof boxes containing ice packs to ensure the freshness and taken to lab.

### Total protein and Enzymatic antioxidant activity estimation

Approximately 0.5mg of fresh leaves were ground in 50mM Sodium Phosphate buffer (pH 7.8) containing 1% polyvinylpyrrolidone (PVP), the homogenate so obtained was centrifuged at 12000g for 15 minutes. The supernatant was diluted to 20ml for protein estimation and enzymatic assays. The protein content was assayed by Lowry’s method using Bovine Serum albumin as standard.

### Superoxide dismutase

Superoxide dismutase activity was assayed by the method described by **Misra and Fridovich (1972)**. Inhibition of Nitroblue tetrazolium reduction by the enzyme was observed using spectrophotometer. The assay volume (3ml) contained 50mM potassium phosphate buffer, 45μM Methionine, 84μM NBT, 5.3μM Riboflavin and 20μM potassium cyanide. Enzyme extract (0.3ml) was added to the experimental tube and all the test tubes were incubated at 25°C Further all the tubes were exposed to fluorescent light for 10 min. The reduced NBT was observed in spectrophotometer at 600nm. One unit of enzyme activity was defined as amount of enzyme required for 50% inhibition of the reduction of NBT and it was expressed as Units mg^−1^ protein.

### Catalase

Catalase activity was assayed by using the method described by **Luck (1974)**. The decomposition of hydrogen peroxide by catalase lead to decrease in absorbance which is read in spectrophotometer (240nm). The enzyme extract (0.2ml) in phosphate buffer (0.67M, pH 7) was observed at 240nm as control and the experimental test tube was added with Hydrogen peroxide. The time required (Δt) for decrease in absorbance from 0.45 to 0.40 was recorded. The time so noted was used for calculation of catalase enzyme activity. The values were expressed as Units mg^−1^ protein.

### Peroxidase

Peroxidase activity was assayed by the method described by **Putter (1974)**. Guaiacol (20mM was used as a substrate and its reaction with hydrogen peroxide (12.3mM) is catalyzed by peroxidase. 1gm of plant tissue was crushed with 3ml of 0.1M phosphate buffer and centrifuged at 18000g for 15min. The supernatant was used as enzyme extract for assay. Phosphate buffer solution (3ml) was added with 0.05ml of Guaiacol, 0.03ml of H_2_O_2_ and 0.1ml of enzyme. The solution was mixed well and left at room temperature for 1min. The absorbance of assay mixture was observed in a spectrophotometer at 436nm and the time (Δt) to increase the absorbance by 0.1 was noted. The enzyme activity was expressed as Units L^−1^.

### Polyphenol oxidase

The polyphenol oxidase activity was assayed by the method of **Esterbauer et al., (1977)**. The change in absorbance due to oxidation of Catechol was recoded at 495nm. Enzyme extract (0.2ml) was mixed with the reaction mixture containing 2.5ml of 0.1M phosphate buffer and 0.3ml of 0.01M Catechol solution. Change in absorbance was recorded in a spectrophotometer at 495nm at an interval of 30sec for 5min. One unit of catechol oxidase is defined as the amount of enzyme that transforms 1μm of dihydrophenol to 1μm of quinine per minute under the assay conditions.

### Ascorbate peroxidase

Ascorbate peroxidase activity was measured using the method described by **Nakano and Asada (1981)**. Reaction mixture (3ml) containing potassium phosphate buffer (50mM, pH 7), ascorbic acid (5mM), and EDTA (1mM) was added with 0.1ml of enzyme extract. H_2_O_2_ (1mM) was added to initiate the reaction. Decrease in absorbance was recorded in a spectrophotometer at 290nm and enzyme activity was calculated by using the extinction coefficient of ascorbate (2.8mM^−1^ cm^−1^). The enzymatic activity can be expressed as U ml^−1^ of enzyme extract.

### Glutathione Reductase

Glutathione reductase activity was assayed as per the method described by **David and Richard (1983)**. The decrease in absorbance due to reduction of oxidized glutathione by the enzyme was observed in a spectrophotometer at 340nm. Enzyme extract (0.1ml) was mixed with 1ml of potassium buffer (0.12 M, pH 7.2), 0.1 ml of EDTA, 0.1ml of sodium azide and 0.1 ml of oxidized glutathione. The volume was made up to 2 ml with water. This mixture was kept at room temperature for 3 minutes and 0.1 ml of NADPH was added to initiate the reaction. The absorbance at 340nm was recorded at intervals of 15 seconds for 2 minutes.

### Statistical analysis

All the assays were performed in triplicates (n=3) and the data shown as mean±SD. The data so obtained was subjected to analysis of variance (ANOVA) for significant variance, Correlation using the Microsoft Excel, 2016. All the curves were also made in Microsoft excel, 2016.

## Results

Higher altitudes have effect on the protein content and enzyme activities. The protein content has a significant (p<0.05) rise with increasing altitude (**Table1, Fig. 1A**). Activity for Catalase and Peroxidase was observed highest in Joshimath population (1986m), for Superoxide dismutase and Ascorbate peroxidase in Pipalkoti population (1339m) while Polyphenol and Glutathione reductase in Gopeshwar (1488m) and Srinagar (606m) population respectively. Negative correlation was observed in the SOD activity (r=−0.321) with the altitude, strong positive correlation was observed in POD (r=0.984), CAT (r=0.932) and GR (r=0.963) activity while weak positive correlation was reported for PPO (r=0.552) and APX (r=0.670) (**Table 2**).

**Table 1.**
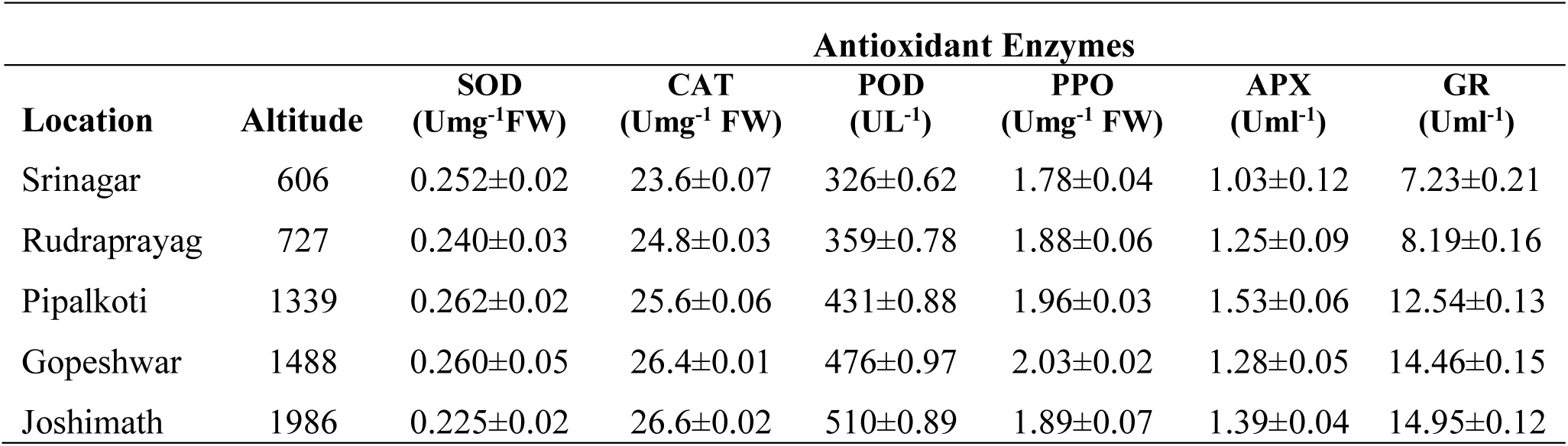
Antioxidant activity of enzymes at different altitudes in Coleus forskohlii

**Table 2.**
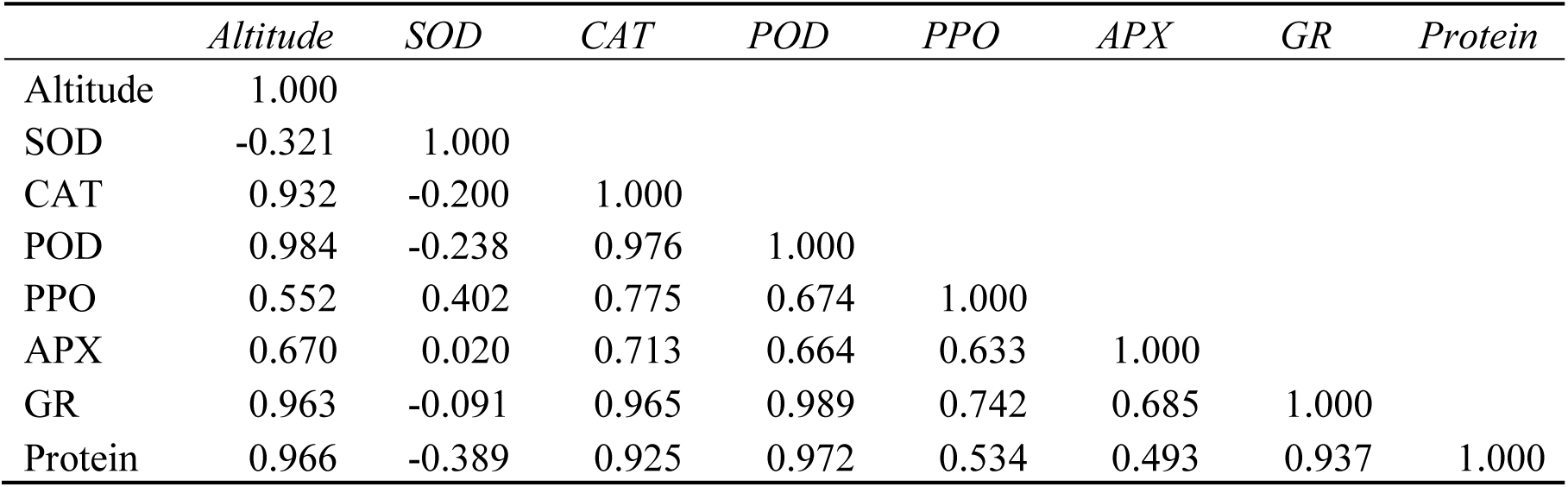
Correlation between antioxidant enzyme and altitude

|  | Altitude | SOD | CAT | POD | PPO | APX | GR | Protein |
| --- | --- | --- | --- | --- | --- | --- | --- | --- |
| Altitude | 1.000 |  |  |  |  |  |  |  |
| SOD | -0.321 | 1.000 |  |  |  |  |  |  |
| CAT | 0.932 | -0.200 | 1.000 |  |  |  |  |  |
| POD | 0.984 | -0.238 | 0.976 | 1.000 |  |  |  |  |
| PPO | 0.552 | 0.402 | 0.775 | 0.674 | 1.000 |  |  |  |
| APX | 0.670 | 0.020 | 0.713 | 0.664 | 0.633 | 1.000 |  |  |
| GR | 0.963 | -0.091 | 0.965 | 0.989 | 0.742 | 0.685 | 1.000 |  |
| Protein | 0.966 | -0.389 | 0.925 | 0.972 | 0.534 | 0.493 | 0.937 | 1.000 |

**Fig. 1.**
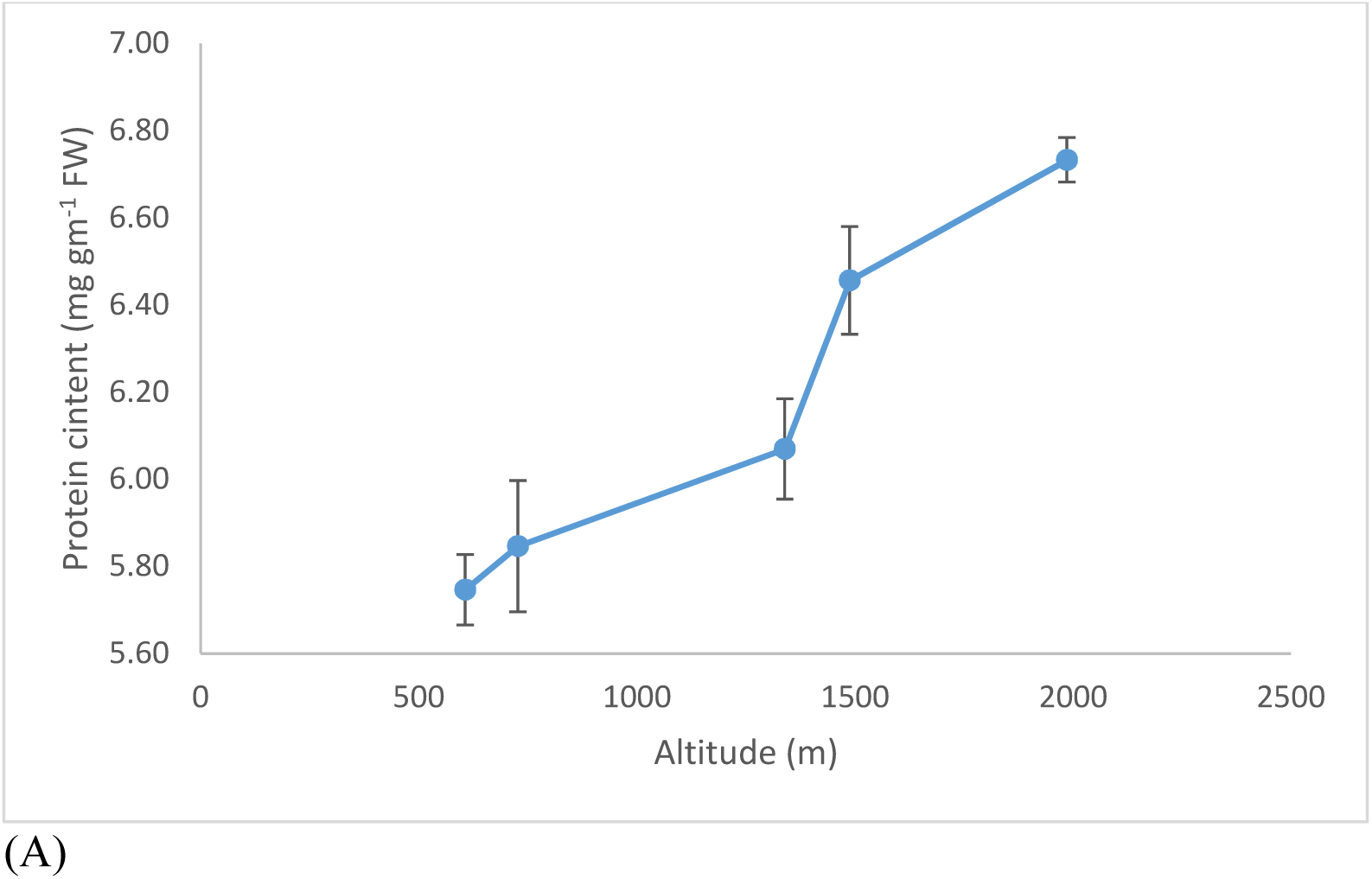
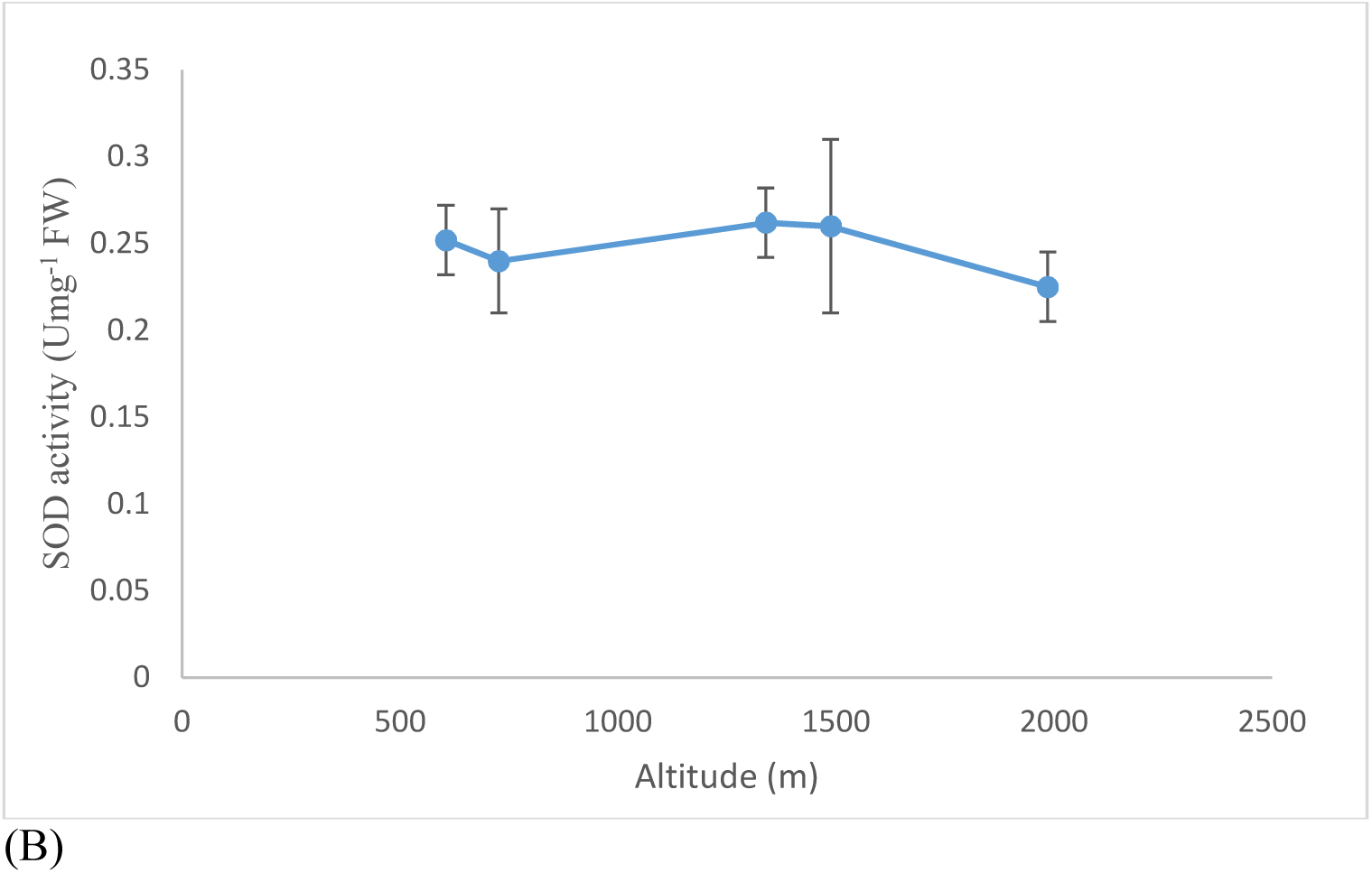
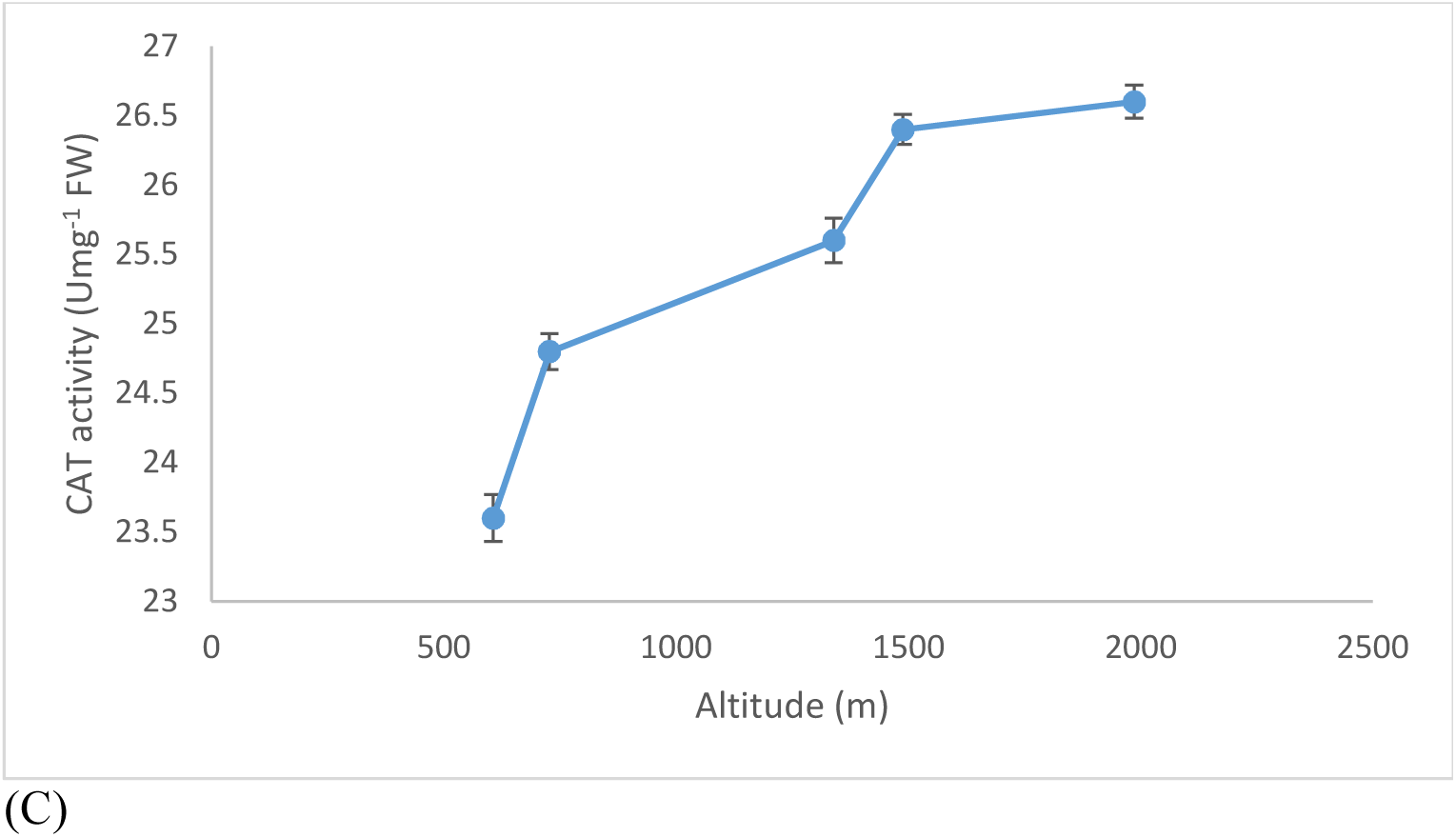
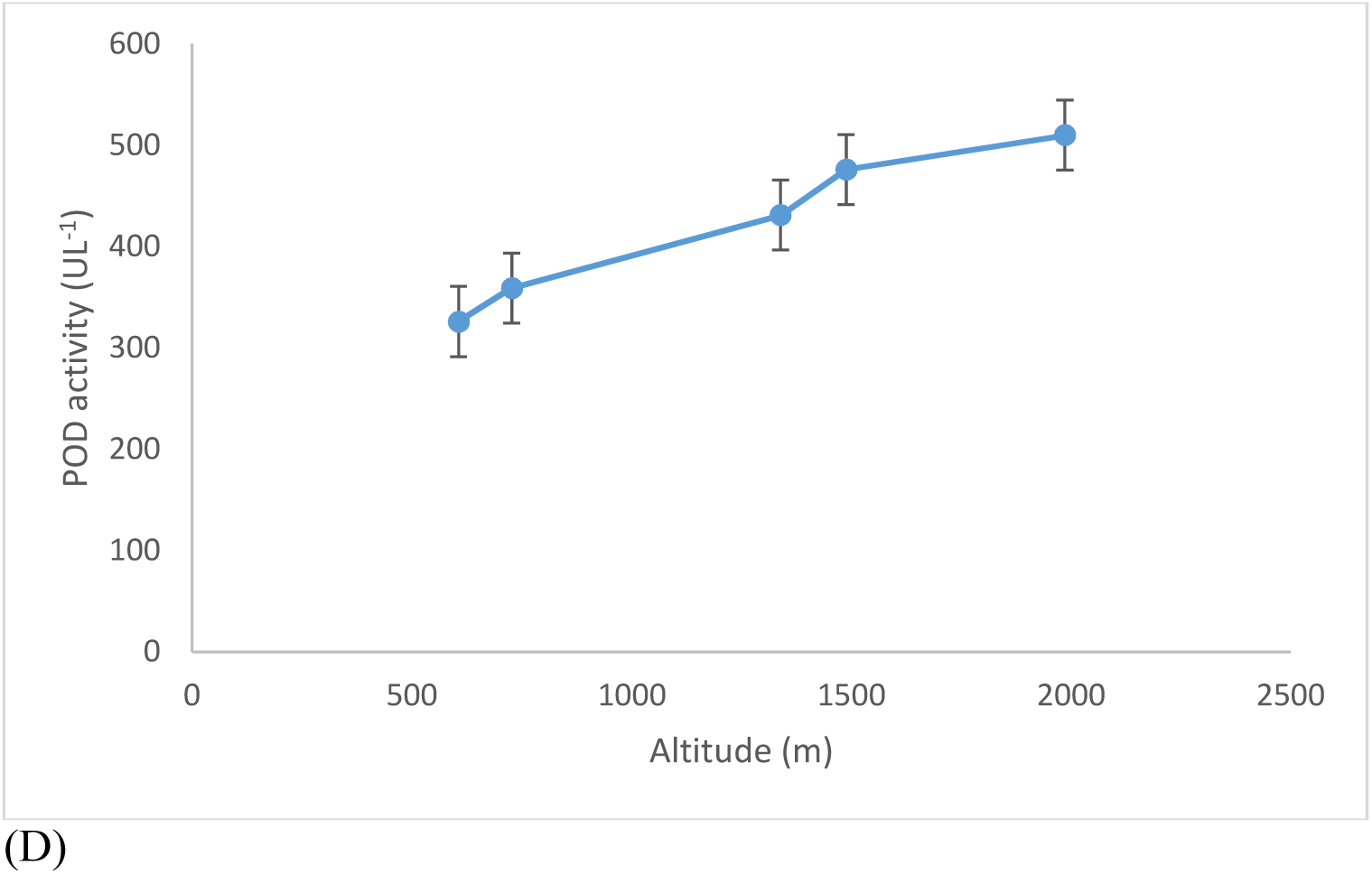
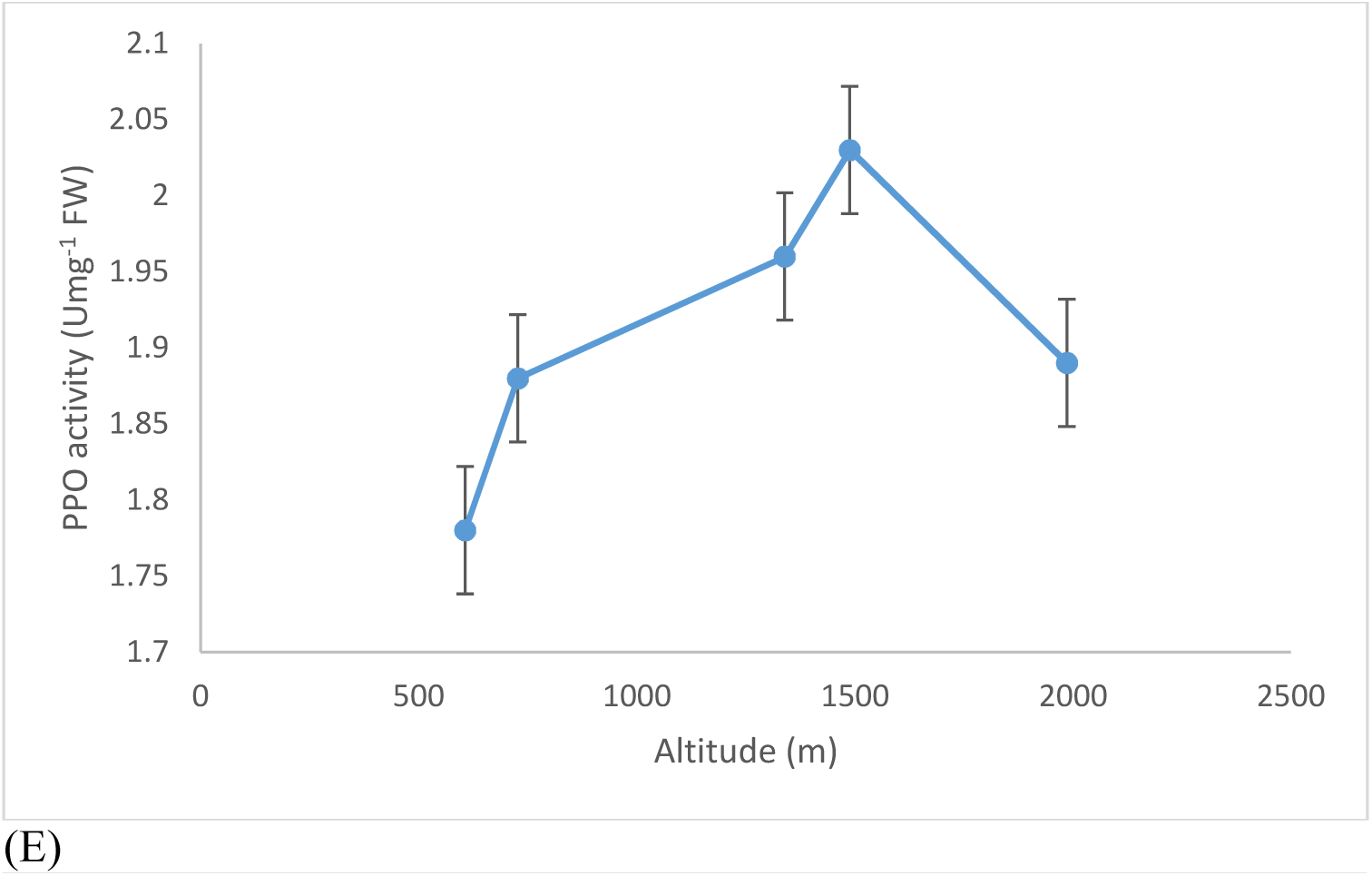
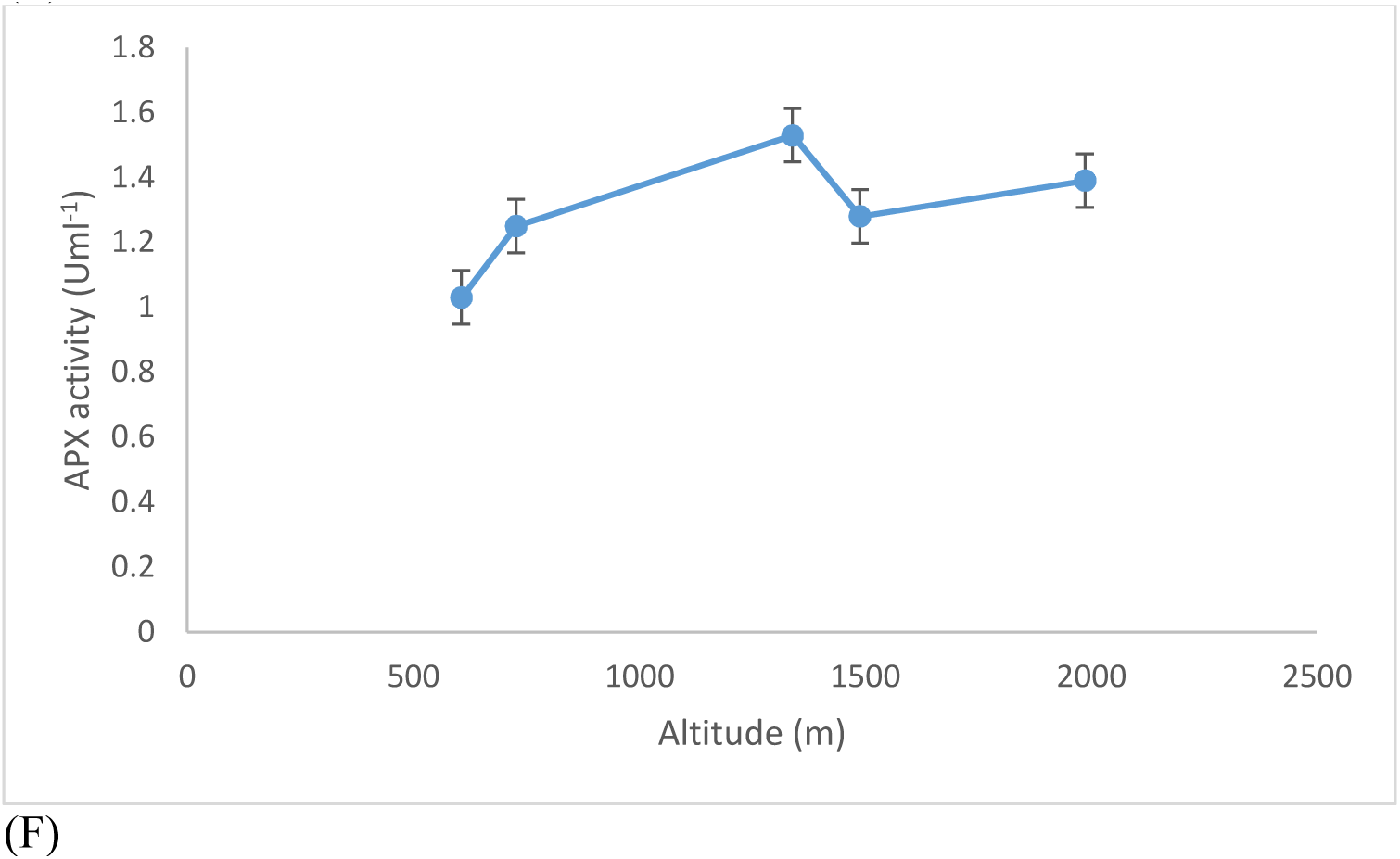
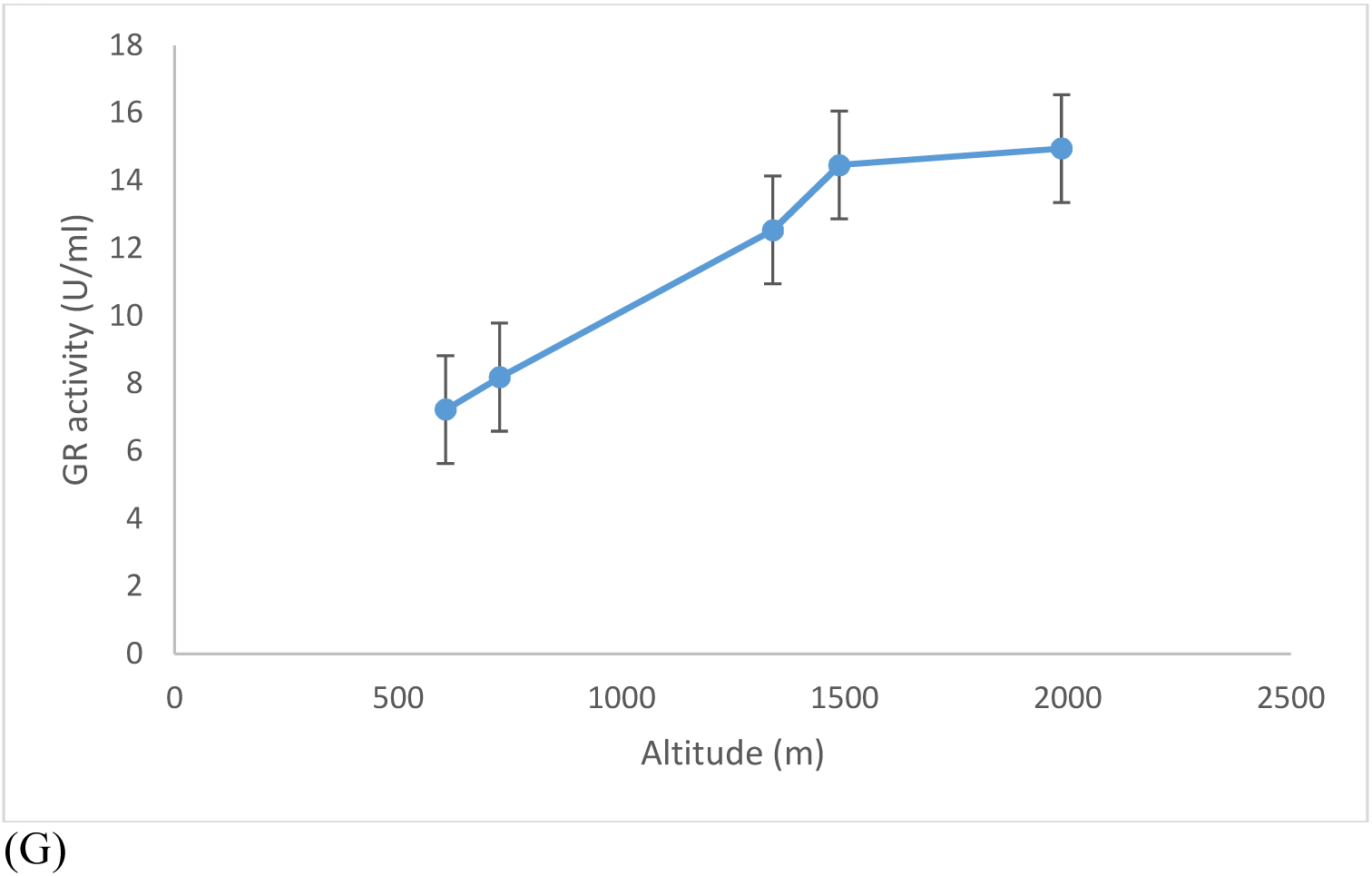
Pattern of protein content and enzymatic activities across altitudes(A) Protein content, (B) SOD, (C) CAT, (D) POD, (E) PPO, (F) APX, (G) GR.

Superoxide dismutase activity was found to be least affected by the altitude. Highest activity was observed in the Pipalkoti (0.262±0.02 Umg^−1^ FW) and Gopeshwar population (0.260±0.05 mg^−1^ FW) while a slight drop in the SOD activity was observed in the Joshimath population as compared to other populations (**Fig. 1B**). Catalase activity was also observed to be increasing with the altitude but no significant increase was observed. Highest activity of Catalase was observed in Joshimath population (26.6±0.02 Umg^−1^ FW) (**Fig. 1C**). Activity of Peroxidase reported with a significant increase over the altitudes and highest value was observed in Joshimath population (510±0.89 UL^−1^) while lowest value was observed in Srinagar population (326 Umg^−1^) (**Fig. 1D**). Polyphenol oxidase activity was found highest in Gopeshwar population (2.03±0.02 Umg^−1^FW) which further decreases in Joshimath population (1.89±0.07 Umg^−1^FW) while lowest activity was recorded in Srinagar population (1.78±0.04 Umg^−1^FW) (**Fig. 1E**). Ascorbate peroxidase activity was observed to be increasing initially with the increasing altitude with lowest activity in the Srinagar population (1.03±0.12 Uml^−1^) and highest activity in Pipalkoti population (1.53±0.06 Uml^−1^) the activity decreases afterwards in Gopeshwar (1.28±0.05 Uml^−1^) and Joshimath (1.39±0.04 Umg^−1^) population (**Fig. 1F**). Highest activity of Glutathione reductase was observed in the Joshimath population (14.95±0.12 Uml^−1^) which decreases at lower altitudes and lowest activity was found in Srinagar population (7.23±0.13 Uml^−1^) (**Fig. 1G**).

## Discussion

Plant growth is affected by various factors like temperature, precipitation, light intensity and radiation intensity. With the increasing altitude all these factors also change which affects the plant growth. In order to get adapted to the changing environmental conditions many physiological and biochemical changes occur in the plant. Effect of environmental factors affecting the physiological and biochemical changes have been studied in many plants (**Cui et al., 2018; Ahmad et al., 2016; Aghaei et al., 2009; Qin et al., 2009; Li et al., 2014**).

In the present study the plant samples were collected from five different altitudes to analyze the effect of changing altitude on the activity of antioxidant enzymes. High protein content in the samples of high altitudes can be attributed to the increased cellular metabolism to cope up with oxidative stress as well as to reduce the osmotic potential under stress (**Basu et al., 2007**). In the present study high protein content in *C. forskohlii* was observed in the plant samples of higher altitude. The increase in protein content under stress conditions may raise the functional protein content to ensure cellular metabolism. With the increasing altitude oxidative stress and intensity of UV radiation increases (**Reiger et al., 2008)** that leads to generation of more reactive oxygen species. In order to combat the oxidative damage due to the ROS plats have developed defense system which involves antioxidant enzymes also. SOD catalyzes the conversion of superoxide anion radicals to hydrogen peroxide and acts as a primary ROS scavenger, an increase in SOD activity indicates towards higher scavenging of superoxide anion radicals (**Asada et al., 1999; Foyar et al., 2005**). CAT plays an important role in defense against oxidative stress damage as it catalyzes the decomposition of hydrogen peroxide to water and oxygen. With their ability to scavenge the free radicals these enzymes help in maintaining the redox state in the plant cell. In the present study very little rise in the levels of SOD and CAT indicates towards better adaptability of *C. forskohlii* and the levels of SOD and CAT are enough to scavenge the deteriorating effect of superoxide anions and hydrogen peroxide. Peroxidase does the similar function of scavenging H_2_O_2_ as Catalase by using electron donors. In the present study the peroxidase activity rises concomitantly with the increasing altitude POD can scavenge the ROS in early senescence and induce lipid peroxidation in late senescence. The opposite trends of CAT and POD may be an adaptive adjustment towards the environment. Positive correlation was reported in Polyphenol oxidase and Ascorbate peroxidase activity with the increasing altitude. Higher APX activity at high altitudes may be to scavenge the plants from high H_2_O_2_ level (**Kaur et al., 2016**). High intensity of UV radiation and the low temperature leads to a higher oxidative stress with H_2_O_2_ production at higher altitudes (**Georgieva et al., 2014**). The increasing activity of GR and PPO at higher altitude may be attributed to the defense mechanism of plant by H_2_O_2_ decomposition. The increasing activity of antioxidant enzymes with the increasing altitude have been reported earlier also (**Khan et al., 2016**).

## Conclusion

This study provides supporting evidence for the higher antioxidant activity of *Coleus forskohlii*. Enhanced activity of SOD, CAT, POD, PPO, APX and GR at higher altitudes reflects higher ROS scavenging potential of *C. forkohlii* population from high altitudes. Thus the population of *C. forskohlii* from a higher altitude can be used a source of high antioxidants.

## Acknowledgement

We gratefully acknowledge the financial support from Uttarakhand Biotechnology Council (UCB), Govt of Uttarakhand, UCB/R&D Project/2017-47.

